# A sub+ cortical fMRI-based surface parcellation

**DOI:** 10.1101/2019.12.20.883553

**Authors:** John D. Lewis, Gleb Bezgin, Vladimir S. Fonov, D. Louis Collins, Alan C. Evans

## Abstract

Both the cortex and the subcortical structures are organized into a large number of distinct areas reflecting functional and cytoarchitectonic differences. Mapping these areas is of fundamental importance to neuroscience. A central obstacle to this task is the inaccuracy associated with mapping results from individuals into a common space. The vast individual differences in morphology pose a serious problem for volumetric registration. Surface-based approaches fare substantially better, but have thus far been used only for cortical parcellation. We extend this surface-based approach to include also the subcortical deep gray-matter structures. Using the life-span data from the Enhanced Nathan Klein Institute - Rockland Sample, comprised of data from 590 individuals from 6 to 85 years of age, we generate a functional parcellation of both the cortical and subcortical surfaces. To assess this extended parcellation, we show that our extended functional parcellation provides greater homogeneity of functional connectivity patterns than do arbitrary parcellations matching in the number and size of parcels. We also show that our subcortical parcels align with known subnuclei. Further, we show that this parcellation is appropriate for use with data from other modalities; we generate cortical and subcortical white/gray contrast measures for this same dataset, and draw on the fact that areal differences are evident in the relation of white/gray contrast to age, to sex, to brain volume, and to interactions of these terms; we show that our extended functional parcellation provides an improved fit to the complexity of the life-span changes in the white/gray contrast data compared to arbitrary parcellations matching in the number and size of parcels. We provide our extended functional parcellation for the use of the neuroimaging community.

## 1. Introduction

Cortical and subcortical gray matter, alike, comprise a large number of areas with distinct functional and structural characteristics. A substantial literature is dedicated to how best to identify these areas. Researchers have divided the cortex into regions that are relatively homogenous with respect to cytoarchitecture (Brodmann, 1909; Hirai and Jones, 1989; Amunts et al., 2005; Zilles and Amunts, 2009, 2010); regions based on morphometry (Tzourio-Mazoyer et al., 2002); regions that are relatively homogeneous with respect to connectivity patterns (Klein et al., 2007; Wig et al., 2013; Gordon et al., 2014); and regions that are relatively homogenous across multiple modalities (Glasser et al., 2016). Each of these approaches has advantages and limitations. One of the most serious limitations, and one that is shared across these parcellation schemes, is the difficulty of mapping the individual results to a common space. Many of these parcellation schemes have relied on volumetric registration techniques to do this. But the normal individual anatomical variability in brain morphology is problematic, particularly when the age-range of the data is large, when both males and females are included, or when abnormalities are present. Moreover, to overcome the failings of volume-based registration methods, the data are often blurred before assessing correlation structure; but this blurs data across tissue classes, and will have very different effects in cortical areas with narrow sulci than in cortical areas with wide sulci, or in gray-matter structures of different sizes; and it blurs areal boundaries. To better deal with the limitations of volume based registration methods, some recent parcellation methods have turned to surface-based registration approaches. Surface-based approaches are significantly more accurate in mapping cortical regions to a common space (Fischl et al., 2007; Lyttelton et al., 2007; Anticevic et al., 2008; Klein et al., 2010). However, to the best of our knowledge, surface-based parcellation approaches have been restricted to the cortex, with subcortical parcellations remaining at the voxel level. But, some of the issues with registration that a surface-based analysis overcomes, e.g. differences in the size and shape of the ventricles, are also issues for registration of the subcortical structures; perhaps even more so.

Accurate registration of the subcortical deep gray-matter structures is notoriously difficult, and inaccurate registration in structures comprised of numerous sub-nuclei will confound parcellation methods; and for structures that are adjacent to the ventricles, like the thalamus and caudate, such inaccuracies will have an even greater impact. Fortunately, recent developments have allowed surfaces to be fitted to the subcortical structures (*e.g.* Lewis et al., 2019). The creation of the cortical surfaces relies on accurate tissue segmentation; but this approach does not work well for the subcortical structures. A multi-atlas label-fusion approach has been shown to fare far better (Aljabar et al., 2009; Collins and Pruessner, 2010; Lötjönen et al., 2010; Coupé et al., 2011; Pipitone et al., 2014). This approach uses a library of labelled atlases, and registers either all of, or a subset of, these atlases to a target image. The registered labels are then fused via *e.g.* patch-based label fusion to produce labels for the target structure. Lewis et al. (2019) used such a method to label the thalamus, caudate, pallidus, and putamen, and then fitted surfaces to these labels. We draw on those methods to extend surface-based parcellation to include those subcortical deep gray-matter structures, as well as the cortex.

Cohen et al. (2008) observed that rs-fMRI connectivity patterns show sharp transitions, which potentially identify areal boundaries. It has since been shown that such transitions correspond to boundaries defined by patterns of cyto- and myeloarchitectonics (Wig et al., 2014; Gordon et al., 2014; Glasser et al., 2016). Gordon et al. (2014) mapped rs-fMRI data to the cortical surface, and built on Cohen et al.’s work to produce all connectivity-based areal boundaries on the cortical surface, and based on these, a cortical parcellation. But Gordon et al. (2014) represented the subcortical deep gray-matter structures in volumetric space. We draw on the methods of Lewis et al. (2019) to extend Gordon et al.’s work to include surface-based representations of the subcortical deep gray-matter structures as well as the cortex. We refer to this cortical+subcortical rs-fMRI connectivity-based parcellation as the Bezgin-Lewis extended Gordon (BLeG) parcellation.

Gordon et al. (2014) assessed their parcellation in multiple ways, including in terms of the homogeneity of the functional connectivity patterns within the parcels in comparison to the same for random variants of that parcellation, as well as the fit of their parcellation to the existing data on cytoarchitectonic boundaries. We extend this assessment to the subcortical structures, and further extend it by drawing on the fact that brain regions show differential relations to age, sex, brain volume, etc. for a variety of morphometric measures (Raz et al., 1997; Goldstein et al., 2001; Allen et al., 2005; Raz et al., 2005; Raz and Rodrigue, 2006; Sowell et al., 2006; Kennedy et al., 2009; Storsve et al., 2014); particularly for subcortical structures (Østby et al., 2009; Goddings et al., 2014). The white/gray contrast measure of Lewis et al. (2018; 2019) provided the basis for a test of this sort of expected within-parcel homogeneity for our cortical+subcortical parcellation. This measure is a ratio of the intensities in the white matter and the adjacent gray matter. Thus it, in part, reflects the integrity of the white matter adjacent to the gray matter, and, in part, the cellular complexity of the gray matter and the degree of myelination within it, including from invading myelinated fibers. And critically, this measure can be produced for cortex and all subcortical structures. Here, the linear model that provided the best fit to the white/gray contrast measures at each vertex was determined for life-span data, as well as the complexity of that model; the homogeneity of model complexity was then computed for every parcel of our extended functional parcellation, BLeG, and the mean of this parcel-wise homogeneity of model complexity provided a measure of how well the overall parcellation aligned with the white/gray contrast data over the life-span. This test was repeated for parcellations based on arbitrary divisions of the surface meshes matching in the number and size of parcels. We show that our extended functional parcellation, BLeG, provides an improved fit to the complexity of the life-span changes in the white/gray contrast data compared to these random parcellations.

## 2. Materials and Methods

### 2.1. Data

The data used here were from the publicly available Enhanced Nathan-Klein Institute - Rockland Sample (Nooner et al., 2012) — commonly known as the NKI-RS data. Data collection received ethics approval through both the Nathan Klein Institute and Montclair State University. Written informed consent was obtained from all participants, and in the case of minors, also from their legal guardians. All imaging data were acquired from the same scanner (Siemens Magnetom TrioTim, 3.0 T). T1-weighted images were acquired with an MPRAGE sequence (TR = 1900 ms; TE = 2.52 ms; voxel size = 1 mm isotropic). Resting state fMRI data were acquired in multiple ways for each subject, varying in temporal and spatial resolution. We utilized the high spatial resolution multiplexed data (TR = 1400 ms; TE = 30 ms; voxel size = 2 mm isotropic). We included all subjects for which there were both usable T1-weighted data and fMRI data. There were 590 such individuals, ranging from 6 to 85 years of age. 64% of these were female.

### 2.2. Data processing

The T1-weighted volumes were processed to derive surfaces onto which the fMRI data could be projected, and from which surface-based measures of white/gray contrast could be computed. These various surfaces are derived from the surfaces that lie at the gray-white boundary and the gray-CSF boundary; the processing that produces these surfaces is described next. The derivation of the surfaces onto which the fMRI is projected is described in section 2.2.1.3 together with a description of the fMRI processing. The derivation of the surfaces from which the measures of white/gray contrast are computed is described in section 2.3.3.1 together with a description of how those measures are computed.

#### 2.2.1. Surface extraction

##### 2.2.1.1. Cortical surface extraction

The T1-weighted volumes were denoised (Manjón et al., 2010) and then processed with CIVET (version 2.1; 2016), a fully automated structural image analysis pipeline developed at the Montreal Neurological Institute^1^. CIVET corrects intensity non-uniformities using N3 (Sled et al., 1998); aligns the input volumes to the Talairach-like ICBM-152-nl template (Collins et al., 1994); classifies the image into white matter, gray matter, cerebrospinal fluid, and background (Zijdenbos et al., 2002; Tohka et al., 2004); extracts the white-matter and pial surfaces (Kim et al., 2005); and maps these to a common surface template (Lyttelton et al., 2007).

##### 2.2.1.2. Subcortical surface extraction

Subcortical segmentation into left and right caudate, putamen, globus pallidus, and thalamus was done using a label-fusion-based labeling technique based on Coupé et al. (2011) and further developed by Weier et al. (2014) and by Lewis et al. (2019). The approach uses a population-specific template library. The library was constructed by clustering (as described in Lewis et al., 2019) the deformation fields from the non-linear transforms produced by CIVET, and using the central-most subject of each cluster to construct the entries in the template library. The number of clusters was specified as the square of the natural log of the number of subjects. To create the library entry for a cluster, the non-linear transform for the central-most subject is inverted and used to warp the ICBM-152-nl template together with the sub-cortical segmentation defined on it; this pair is then added to the template library. The template library is thus a set of warped copies of the ICBM-152-nl template together with their correspondingly warped segmentations, and represents the range of deformations found in the population. Once the template library has been created, each subject in the population is non-linearly registered to the *n* closest templates in the library (here, *n* = 7), and the resulting transforms are used to warp their corresponding segmentations to the subject; the final labelling is then established via patch-based label fusion.

Once the subcortical structures for a subject are labeled, surfaces defined on the ICBM-152-nl template are fitted to these labels. These model surfaces are warped to each individual based on the transforms derived from the label-fusion-based labeling stage, and then adjusted to the final labels by moving vertices along a distance map created for each label. The surfaces for each structure are then registered to their corresponding common surface template to ensure cross-subject vertex correspondence, as per the cortical surfaces.

##### 2.2.1.3. rs-fMRI surfaces

Based on these surfaces at the gray-white boundary and gray-CSF boundary, several additional surfaces were created to allow for the surface-based fMRI analysis. The fMRI data were preprocessed (as described in section 2.2.2.1) and then projected onto the cortical midsurface, a surface falling half way between the surface at the cortical gray-white boundary and the surface at the gray-CSF boundary, and onto surfaces 2 mm inside of the surfaces at the white-gray boundary of the subcortical structures. These choices avoid partial volume effects, to the extent possible. These surfaces are shown in Figure 1.

**Figure 1:**
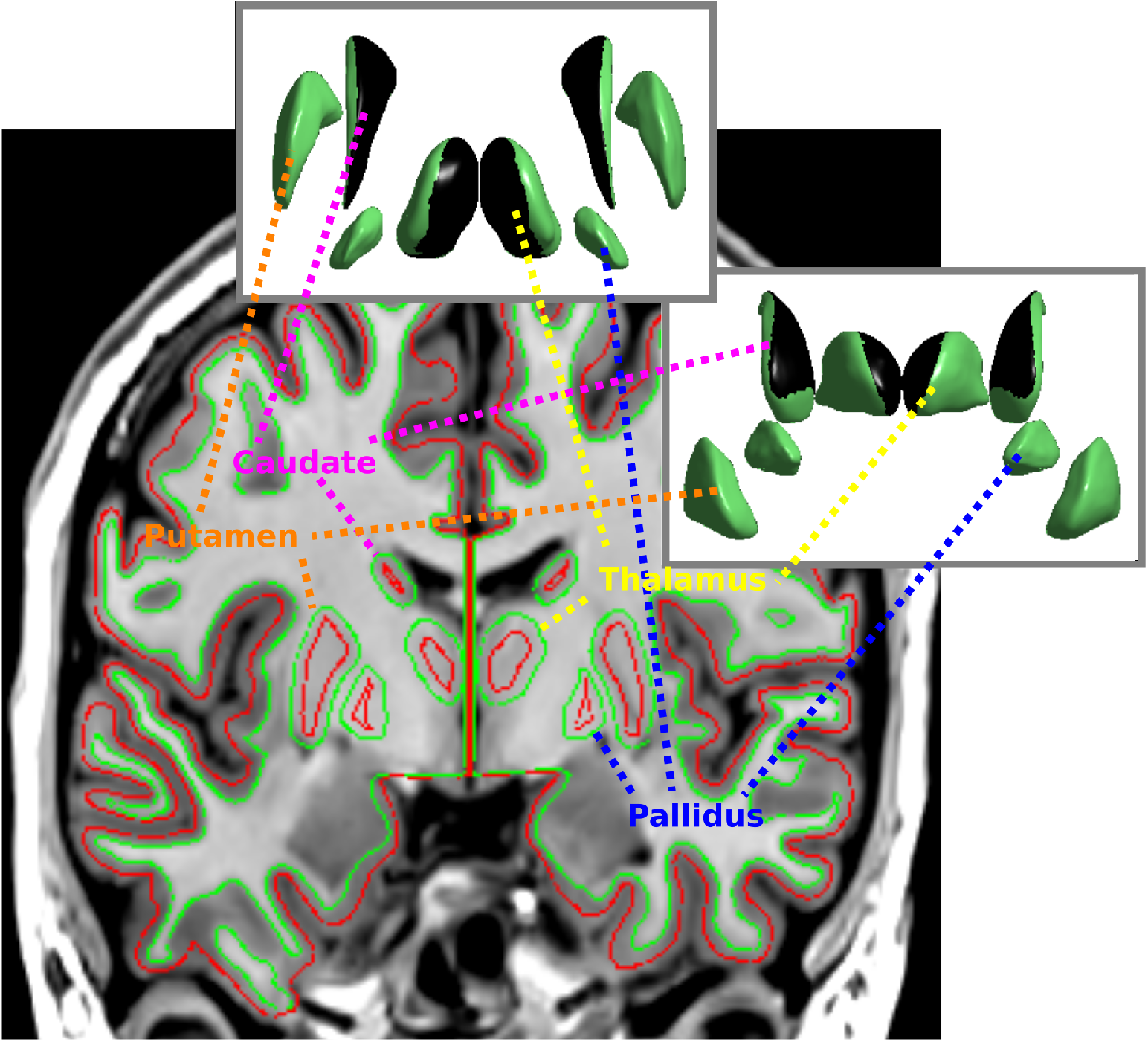
An example of the surfaces that the preprocessed fMRI time-series data are projected onto. The coronal slice shows the cortical midsurface in red, as well as the supra-white surfaces within the subcortical gray matter; the fMRI data are projected onto these surfaces. For reference, the surfaces at the gray-white boundary are shown in green. The insets show the top and front views of the subcortical surfaces, with the areas of the thalamus and caudate that are adjacent to the ventricles shown in black. Note that the spatial aspect of the subcortical structures has been manipulated to provide a view of each structure in each orientation.

To create the surfaces 2 mm inside of the surfaces at the gray-white boundary of the subcortical structures, a distance map was created from those surfaces, smoothed with a Gaussian kernel, and used to create a gradient vector field. The subcortical white surfaces were then moved 2 mm along this vector field into the subcortical gray matter to produce the sub-cortical supra-white surfaces. The procedure is described in detail in Lewis et al. (2019).

#### 2.2.2. rs-fMRI processing

##### 2.2.2.1. rs-fMRI preprocessing

The rs-fMRI data were minimally preprocessed before loading the data onto the surfaces. First, slice-timing correction was applied with FSL’s *slice-timer* (Smith et al., 2004), with the timings for each slice in the multi-band acquisition provided. FSL’s motion correction algorithm, *mcflirt* (Jenkinson et al., 2002) was then used to register all volumes to a reference functional volume. As well as producing the motion-corrected fMRI data, this procedure identified motion-contaminated volumes by frame-by-frame displacement (Power et al., 2012). The fMRI images were then corrected for the distortions associated with echo-planar imaging using large deformation diffeomorphic metric mapping (Miller et al., 2005) based on the T1-weighted volume, with the deformation restricted to the phase encoding direction of the fMRI acquisition. The surfaces described above were then transformed to overlay the distortion-corrected rs-fMRI, and the rs-fMRI data were read onto them, *i.e.* onto the cortical midsurface and the subcortical supra-white surfaces.

##### 2.2.2.2. Connectivity-based parcellation

The connectivity-based parcellation of the rs-fMRI is based on the methods described in Gordon et al. (2014), which builds on the work of Cohen et al. (2008). Cohen et al. (2008) observed that rs-fMRI connectivity patterns show sharp transitions, which potentially identify areal boundaries. Gordon et al. (2014) extended Cohen et al.’s work to produce all cortical connectivity-based areal boundaries, and based on these, a parcellation of the cortex. We further extend this work by including surfaces for the caudate, globus pallidus, putamen, and thalamus, in addition to those for the cortex, producing a parcellation for all of these surfaces. To achieve this, we adapt the code provided by Gordon et al. (2014). We convert the cortical midsurfaces and subcortical supra-white surfaces, with their associated rs-fMRI time courses, to CIFTI, and as per Gordon et al. (2014), for each subject, correlate the time course of each surface vertex (both cortical and subcortical) with that from every other surface vertex. The resultant correlation maps are then transformed using Fisher’s r-to-z transformation. The pairwise correlations between entries in each subject’s correlation map comprise that subject’s similarity matrix. A set of gradient maps that identify positions of abrupt changes in connectivity patterns are then generated by taking the first spatial derivative in each subject’s similarity matrix. The gradient maps are then averaged across subjects, and the average gradient maps subjected to Beucher’s (1979) “watershed by flooding” algorithm to identify potential areal boundaries. The resulting boundary maps are then averaged to yield a map of the frequency with which each vertex is identified as a potential boundary vertex. The resulting boundary map for the NKI-RS data is shown in Figure 2 together with its associated parcellation, derived as described below.

**Figure 2:**
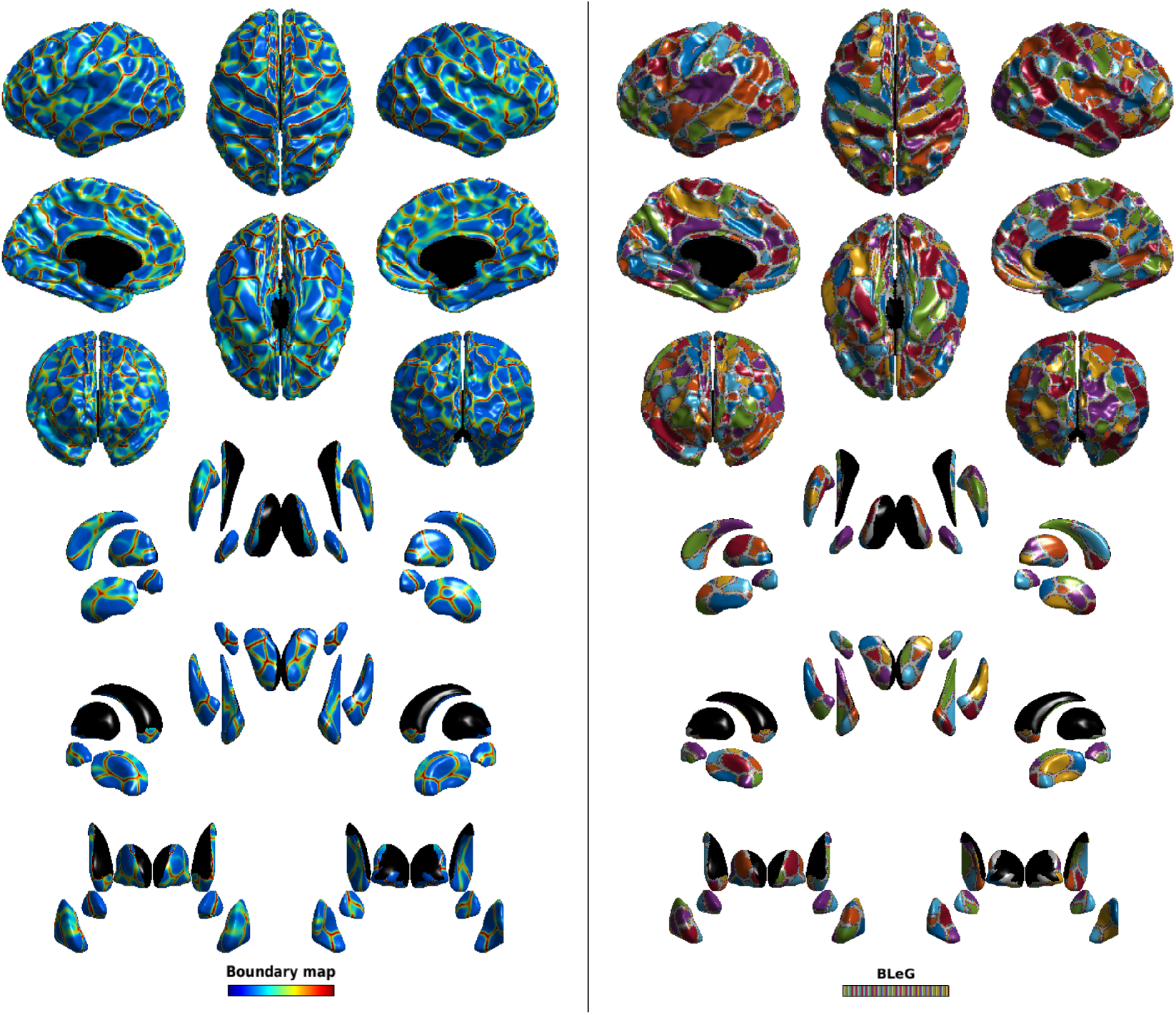
The connectivity-based boundary map (left) and the resulting BLeG parcellation (right). The top half of the figure shows the cortical results; the bottom half shows the subcortical results, with the same set of orientations as the cortical results. Note that the spatial aspect of the subcortical structures has been manipulated to reveal the map of each structure in each orientation; but the position of each structure is nonetheless approximately correct. Notice that the watershed boundaries are generally quite clear and translate directly to the parcel boundaries.

The local minima in the boundary map are seeds for parcel creation. Parcels are expanded outward from these seeds until they either meet another parcel or reach a height threshold on the boundary map. Adjacent parcels are then merged, if they are too similar. Additionally, parcels containing fewer than 30 cortical vertices are merged with the adjacent parcel with the lowest boundary map values separating the two. Lastly, vertices with high boundary map values (defined as the top quartile of values in the boundary map) were eliminatd from parcels, and treated as transition zones between parcels.

### 2.3. Assessing the parcellation

Gordon et al. (2014) assessed their parcellation in multiple ways, *e.g.* in terms of the within-parcel homogeneity of functional connectivity patterns, and by visually comparing parcel alignment with known cytoarchitectonic areas. We extend these assessments to include the subcortical structures, and further, to data from another modality. We assess our parcellation in terms of the within-parcel homogeneity of the complexity of the models required to best-fit white/gray contrast measures (Lewis et al., 2018, 2019). We use white/gray contrast measures because they span both the cortical and the subcortical structures and have been shown to be sensitive to age, sex, and brain size. We determine the linear models that best fit the data at each vertex, and the complexity of those models, and then assess the within-parcel homogeneity of model complexity. The mean of this parcel-wise homogeneity of model complexity provides a measure of how well the overall connectivity-based parcellation aligns with the age, sex, and brain size related differences in white/gray contrast measures. Both assessments of within-parcel homogeneity are done for the BLeG parcellation in comparison to parcellations based on arbitrary divisions of the surface meshes but with the same number of parcels and same mean parcel size as the BLeG parcellation.

#### 2.3.1. Assessment of BLeG with fMRI connectivity patterns

For each parcellation, we determined the parcel-wise homogeneity of the fMRI connectivity patterns. We define the homogeneity of fMRI connectivity for a parcel *p* to be the the mean of the correlations between the connectivity patterns of all pairs of vertices within parcel *p*, across subjects. We define the within-parcel homogeneity of fMRI connectivity for the overall parcellation as the mean of the parcel-wise homogeneity.

We assess the homogeneity of fMRI connectivity for our BLeG parcellation and for parcellations based on arbitrary divisions of the surface meshes. We generated 100 parcellations with random parcels. These were random in terms of the placement of the parcels, but each random parcellation had the same number of parcels in each structure as does the BLeG parcellation, and was constructed such that the mean size of the parcels did not differ significantly from the BLeG parcellation. This is shown in Figure 3. We then compute the mean and standard deviation of the within-parcel homogeneity of fMRI connectivity across the 100 random parcellations, and the relation of the BLeG parcellation to these random parcellations in terms of standard deviations from the mean of the random parcellations.

**Figure 3:**
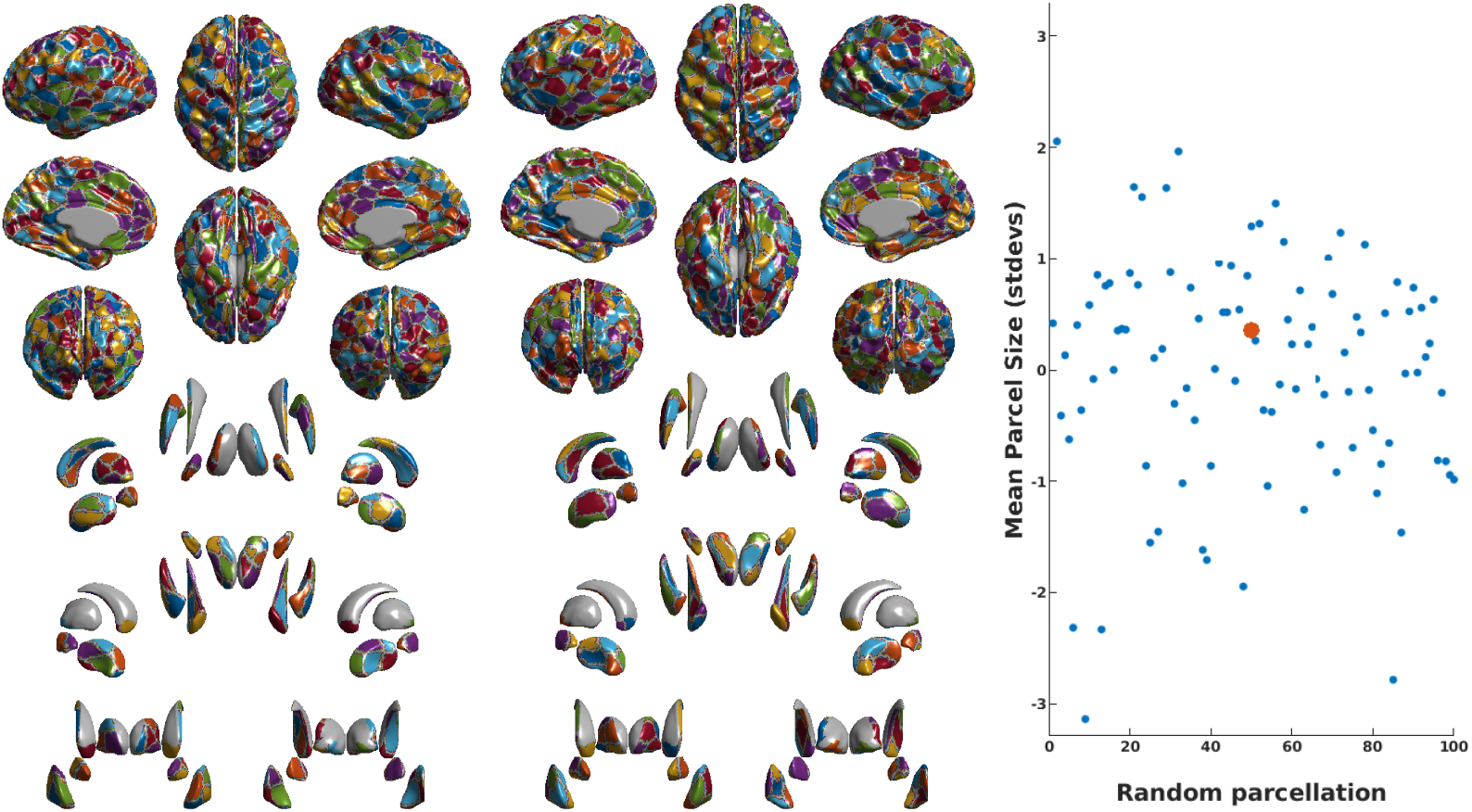
Two examples of random parcellations and a comparison of the mean parcel size of all 100 random parcellations to that of the BLeG parcellation. The mean parcel size of the BLeG parcellation (red dot) is cleary not significantly different from that of the random parcellations (blue dots).

#### 2.3.2. Comparison of BLeG subcortical parcels to known anatomy

Gordon et al. (2014) demonstrated that these methods produce a parcellation that, at least for the cortex, aligns well with known cytoarchitectonic areas. They visually compared several cortical parcels to the probabilistic borders of cortical areas that were mapped by Van Essen et al. (2011) based on cytoarchitectonic maps produced by Fischl et al. (2007). We extend this approach to the subcortical structures; but lacking probabilistic maps for these structures, we rely on pictorial descriptions of the anatomy of the structures. A consensus description is available only for the thalamus; we compare our BLeG parcellation of the thalamus to that.

#### 2.3.3. Assessment of BLeG with white/gray contrast

##### 2.3.3.1. White/gray contrast measurements

In order to generate the white/gray contrast measures, two additional surfaces were created from each of the surfaces at the gray-white boundary: a supra-white surface inside the gray matter, and a sub-white surface inside the white matter. The T1-weighted intensities were then projected onto these surfaces, and at each vertex, the value on the sub-white surface was divided by the value on the supra-white surface. For the cortex, the sub-white surface was placed 1 mm beneath the surface at the inner edge of the cortical gray matter and the supra-white surface was placed 35% of the way from the surface at the gray-white boundary to the surface at the gray-CSF boundary. For the subcortical structures, because of the lesser spatial constraints, the sub-white surfaces were placed 2 mm outside of the surfaces at the gray-white boundaries, and the supra-white surfaces were placed 2 mm inside of the surfaces at the gray-white boundaries. These surfaces are shown in Figure 4. To create the surfaces on either side of the gray-white boundary, a distance map was created from the surfaces at the gray-white boundary (both cortical and subcortical), smoothed with a Gaussian kernel, and used to create a gradient vector field. The cortical white surface was moved 1mm inward along this gradient vector field to produce a sub-white surface, and outward 35% of the distance to the gray surface to produce a supra-white surface. The same procedure produced the contrast measures for the subcortical surfaces, but moving inward and outward 2 mm. This procedure ensures that the subwhite surfaces in regions with thin strands of white matter will not cross, and so will provide the best possible approximation of white matter. This can be seen in Figure 4 between subcortical structures and within thin gyri. Notice, however, that areas of the subwhite surfaces of the caudate and thalamus fall within the ventricles rather than white matter, and thus the white/gray contrast measures in these areas will not be valid and must be masked. This is, of course, also the case for the brain stem and midsagittal cuts.

**Figure 4:**
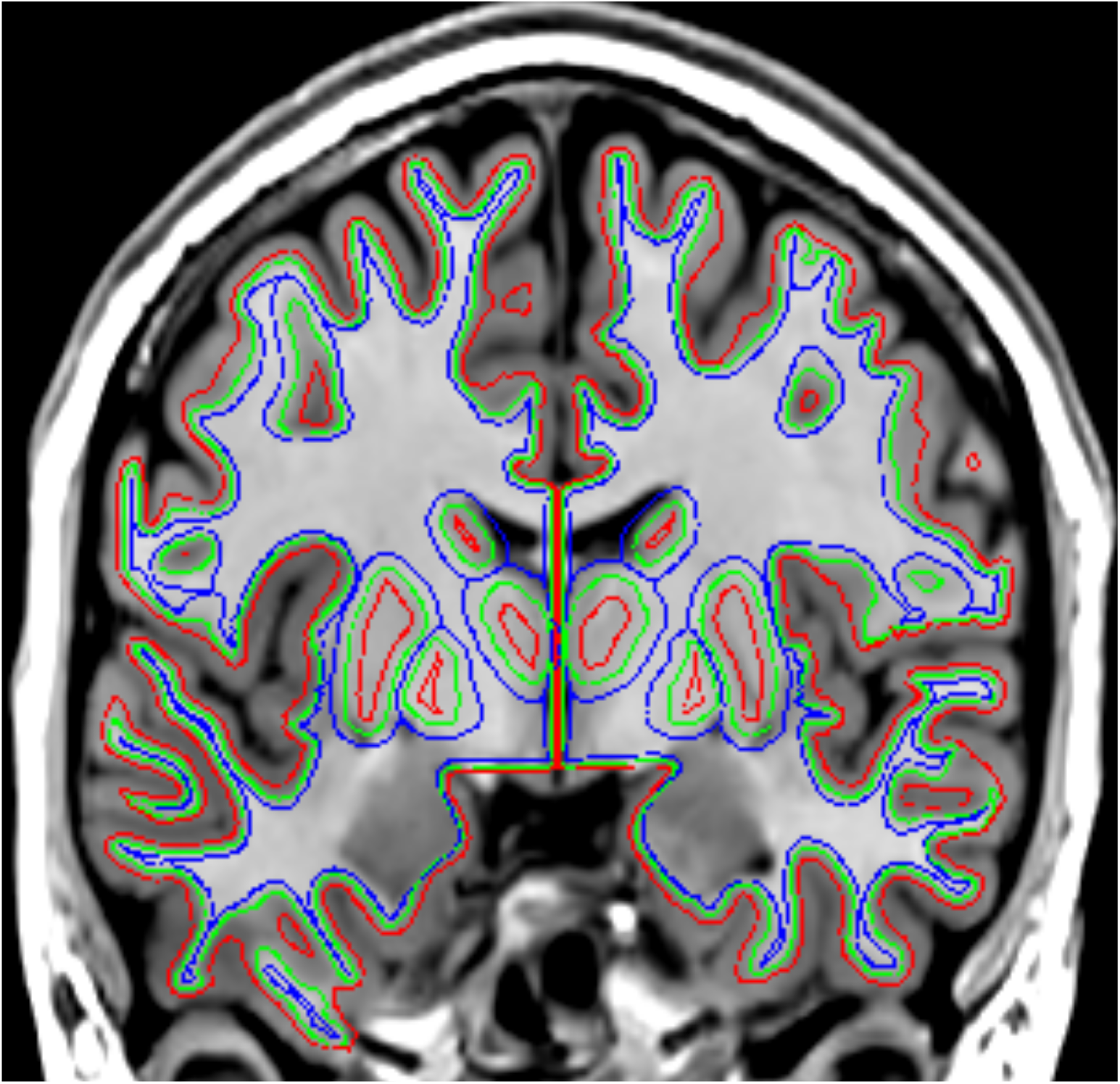
An example of the surfaces used to form the the white/gray contrast measures. The surface at the gray-white boundary is shown in green. Copies of these surfaces were moved into gray matter (red) and white matter (blue) along the gradient vectors of a distance map based on the surface at the gray-white boundary. Gray matter and white matter intensity were then measured at each of the vertices of these derivative surfaces, and the white/gray contrast measure formed as the ratio of white intensity to gray intensity at corresponding vertices of the two surfaces. Note that in areas with thin strands of white matter subwhite surfaces abutt rather than cross. Note also that areas of the subwhite surface of the caudate and thalamus fall in the ventricles; measures in these areas are invalid and must be masked.

The intensity values on the T1-weighted image were sampled at each vertex of both the supra-white surfaces and the sub-white surfaces, smoothed with a 10 mm FWHM blurring kernel to reduce measurement noise without excessively blending measures from different brain regions, and the ratio was formed by dividing the value at each vertex of the sub-white surface by the value at the corresponding vertex of the supra-white surface. The intensity values were sampled in stereotaxic space, with the T1-weighted volume up-sampled to 0.5 mm iso, with no non-uniformity correction or normalization. This avoids, to the extent possible, issues arising from differences in brain size, while leaving the intensity values essentially unchanged.

##### 2.3.3.2. Within-parcel complexity homogeneity of BLeG

At each vertex, we determined the best-fit model to the contrast data from all models comprised of terms including any of ‘AGE’, ‘SEX’, and ‘BRAINVOL’ (total), as well as any of the composite terms ‘AGE^2^’, ‘AGE^3^’, ‘AGE^4^’ and the interaction terms for any of ‘SEX’, ‘BRAINVOL’, and ‘AGE^1‥4^’. The best-fit model was determined by searching all possible models and choosing the one with the lowest value for the Akaike information criterion (AIC) (Akaike, 1976). Each model was evaluated using the SurfStat toolbox^2^. We then determined the complexity of the best-fit model at each vertex. We defined *model complexity* as the sum of the number of terms, with composite terms counted as the number of base terms in the composite term raised to the power 0.5, capturing the intuition that *e.g.* ‘1 + AGE + AGE^2^’ should be more complex than ‘1 + AGE + SEX’, but less complex than ‘1 + AGE + SEX + BRAINVOL’. This yielded the map in Figure 5. For each parcellation, we determined the parcel-wise homogeneity of model complexity. We define the homogeneity of model complexity for a parcel *p* to be 1/(1 + *std*(*mc*_*p*_)), where *mc*_*p*_ is the vector comprised of the model complexity at each vertex within parcel *p*. We define the homogeneity of model complexity of the overall parcellation as the mean of the parcel-wise homogeneity of model complexity. We assess the homogeneity of model complexity for our BLeG parcellation and for the parcellations based on arbitrary divisions of the surface meshes described in section 2.3.1. We then compute the mean and standard deviation of the homogeneity of model complexity across the 100 random parcellations, and the relation of the BLeG parcellation to these random parcellations in terms of standard deviations from the mean of the random parcellations.

**Figure 5:**
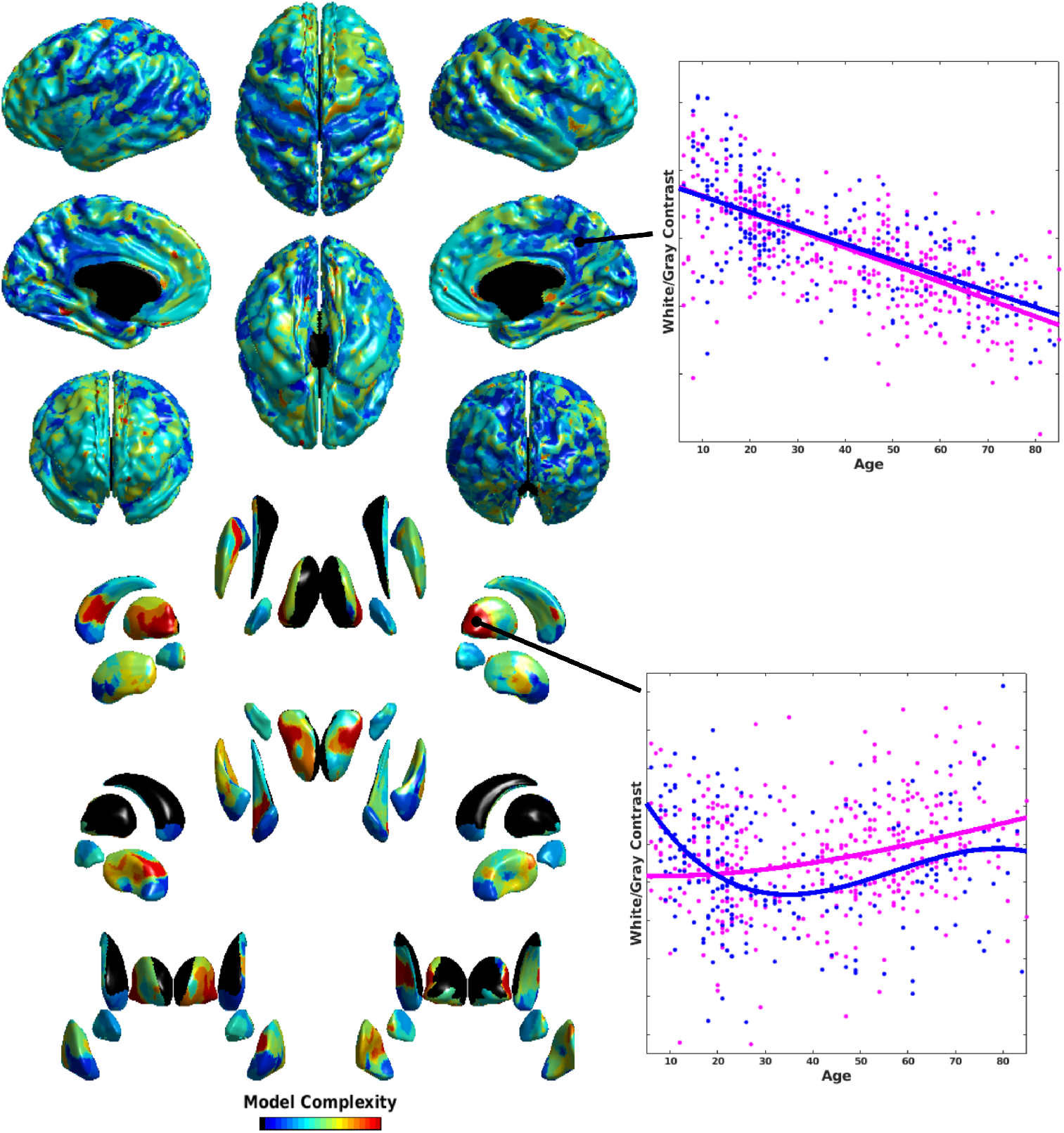
The map of the complexity of the best-fit models at each vertex. Note that the spatial aspect of the subcortical structures has been manipulated to provide a view of each structure in each orientation. Notice that model complexity varies greatly over the cortex, and even more so in the subcortical structures. The scatter plots to the right show the white/gray contrast data plotted against age at the points indicated, with female data shown in magenta and male data shown in blue.

## 3. Results

### 3.1. Assessment of BLeG with fMRI connectivity patterns

The results of the test of the homogeneity of within-parcel fMRI connectivity based patterns for the BLeG parcellation is shown in Figure 6. The BLeG parcellation shows mean within-parcel fMRI connectivity homogeneity more than 27 standard deviations greater than that for the mean of random parcellations with the same number of parcels in each structure and a mean parcel size that is not different from the BLeG parcellation. As shown in the right subfigure of Figure 6, this greater mean within-parcel homogeneity of model complexity corresponds to broadly increased within-parcel homogeneity of fMRI connectivity patterns, with substantially greater increased homogeneity present in subcortical structures. Thus, the BLeG parcellation captures the patterns of functional connectivity to a far greater extent than would be expected from the size of the parcels alone, and this is true for parcels throughout the cortex and the subcortical structures.

**Figure 6:**
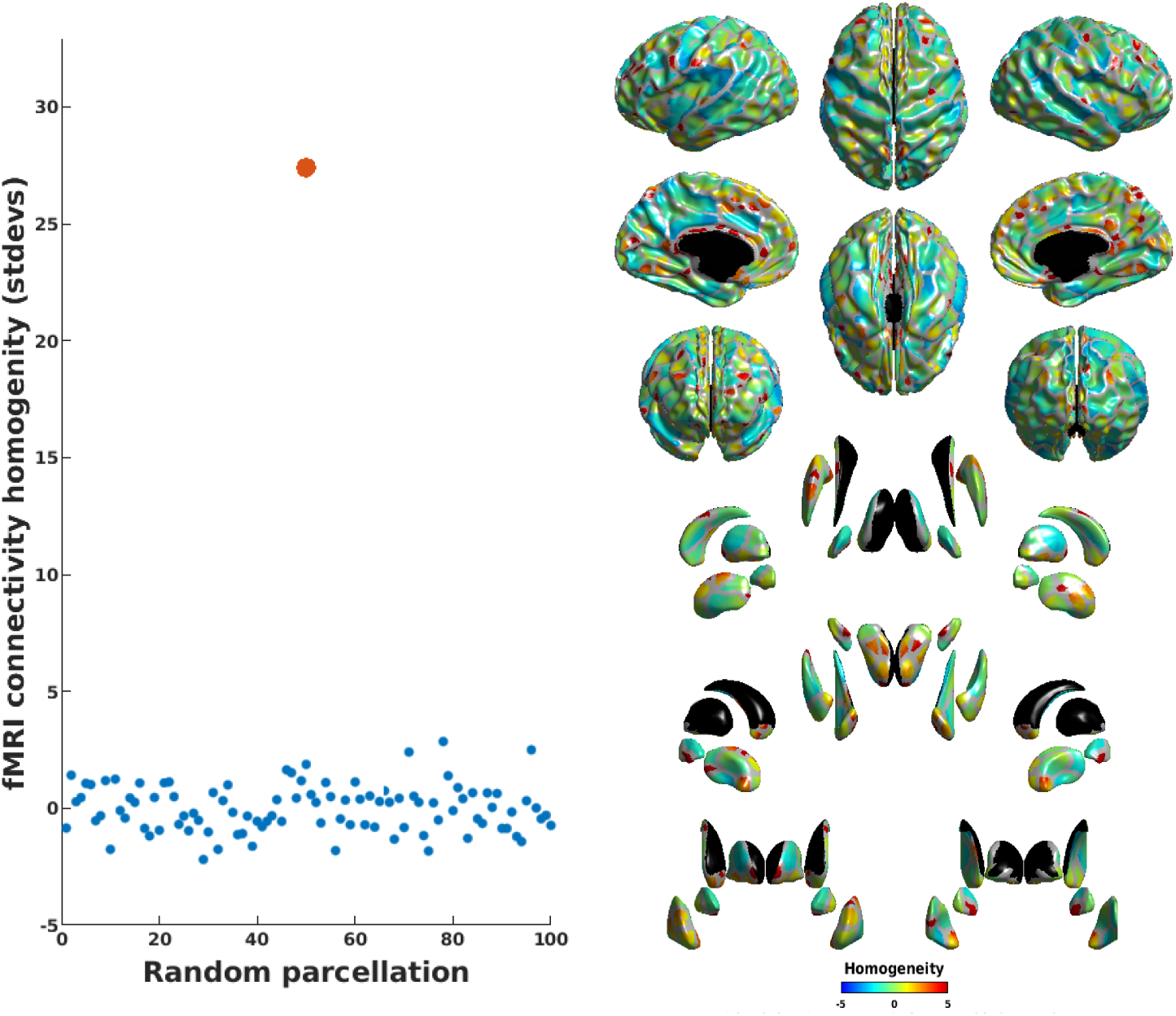
fMRI connectivity homogeneity. The left subfigure shows the comparison of mean within-parcel homogeneity of fMRI connectivity for the BLeG parcellation (red) vs random parcellations (blue). The BLeG parcellation shows mean within-parcel homogeneity more than 27 standard deviations greater than that for the random parcellations. The subfigure on the right shows the pattern of the increased homogeneity in the BLeG parcellation.

### 3.2. Comparison of BLeG subcortical parcels to known anatomy

The comparison of our BLeG parcellation to the pictorial description of the consensus anatomy of the thalamus is shown in Figure 7. Our BLeG parcellation excludes the portion of the thalamus adjacent to the ventricles, which is approximately the region medial to the internal medullary lamina, and so the comparison applies only to the lateral nuclei. This comparison yields a good correspondence. The consensus anatomy shows eight subnuclei internal to the thalamus; each of these can be directly mapped to the eight thalamic BLeG parcels. Note also the symmetry of the left and right thalami; this also agrees with the literature. The lateral and medial geniculate nucleus are not represented in the BLeG parcellation, since the fMRI data were sampled 2mm inside the surface of the thalamus, and so not inside these very small structures that protrude from the thalamus.

**Figure 7:**
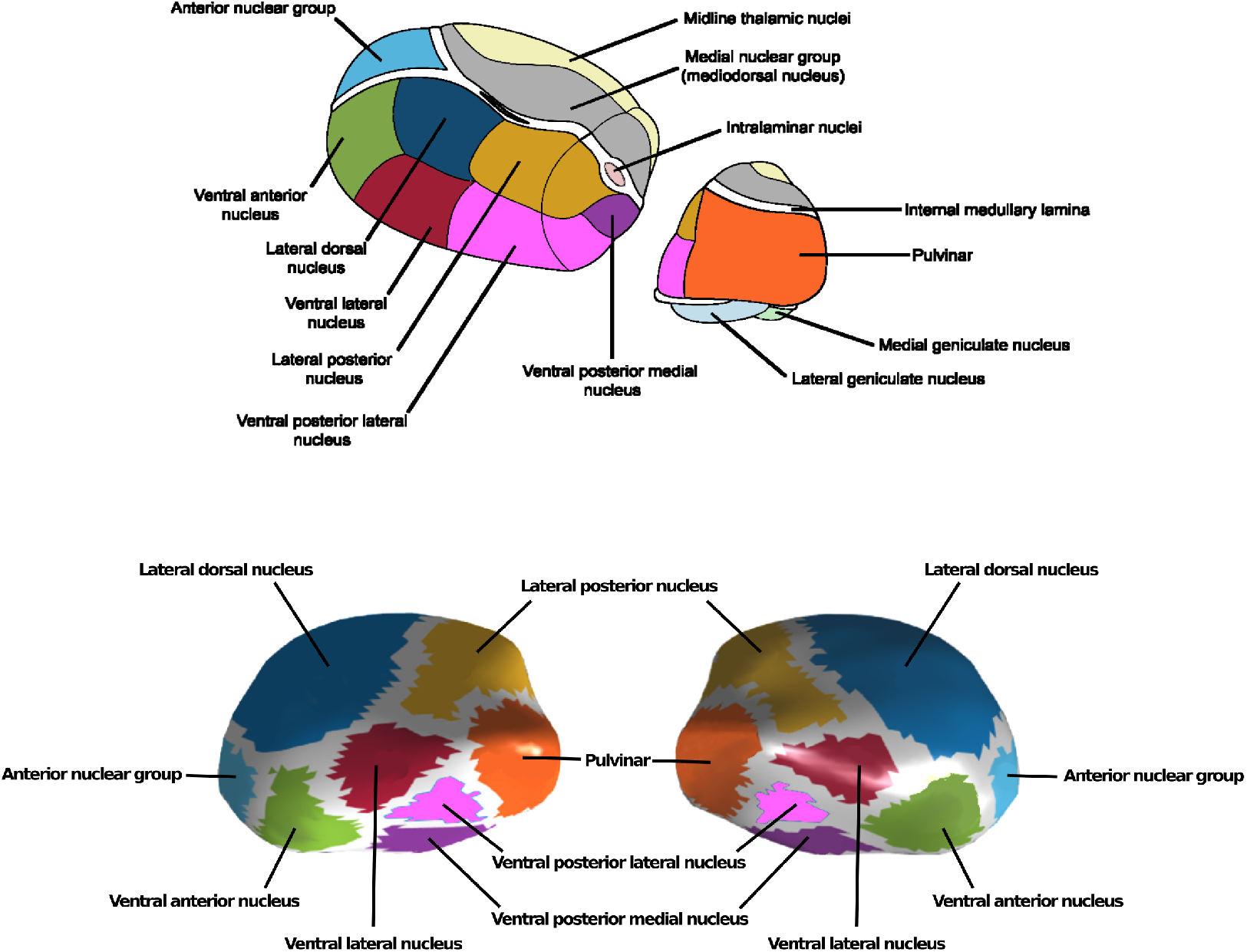
Comparison of the BLeG parcellation of the thalamus to the pictorial description of its consensus anatomy. Note that the pictorial description of the thalamus is viewed from above in order to show the midline thalamic nuclei and the medial nuclear group, whereas our BLeG parcellation masks the portion of the thalamus adjacent to the ventricles, essentially discarding the areas medial to the internal medullary lamina, and so is shown in a lateral view. Comparing the BLeG parcellation to the consensus anatomy for the portion of the thalamus lateral to the internal medullary lamina shows a perfect correspondence. Note that the fMRI data are measured 2mm inside the thalamic surface, and so activity in the lateral and medial geniculate nuclei is not measured.

### 3.3. Assessment of BLeG with white/gray contrast

The results of the test of the alignment of our BLeG parcellation with the map of the model complexity for the white/gray contrast data are shown in Figure 8. The BLeG parcellation shows mean within-parcel model complexity homogeneity almost 10 standard deviations greater than that for the mean of random parcellations with the same number of parcels in each structure and a mean parcel size that is not different from the BLeG parcellation. As shown in the right subfigure of Figure 8, this greater mean within-parcel homogeneity of model complexity corresponds to broadly increased within-parcel model homogeneity, with substantially greater increased homogeneity present in subcortical structures. Thus, the BLeG parcellation captures the patterns of model complexity within the white/gray contrast data to a far greater extent than would be expected from the size of the parcels alone, and this is true for parcels throughout the cortex and the subcortical structures.

**Figure 8:**
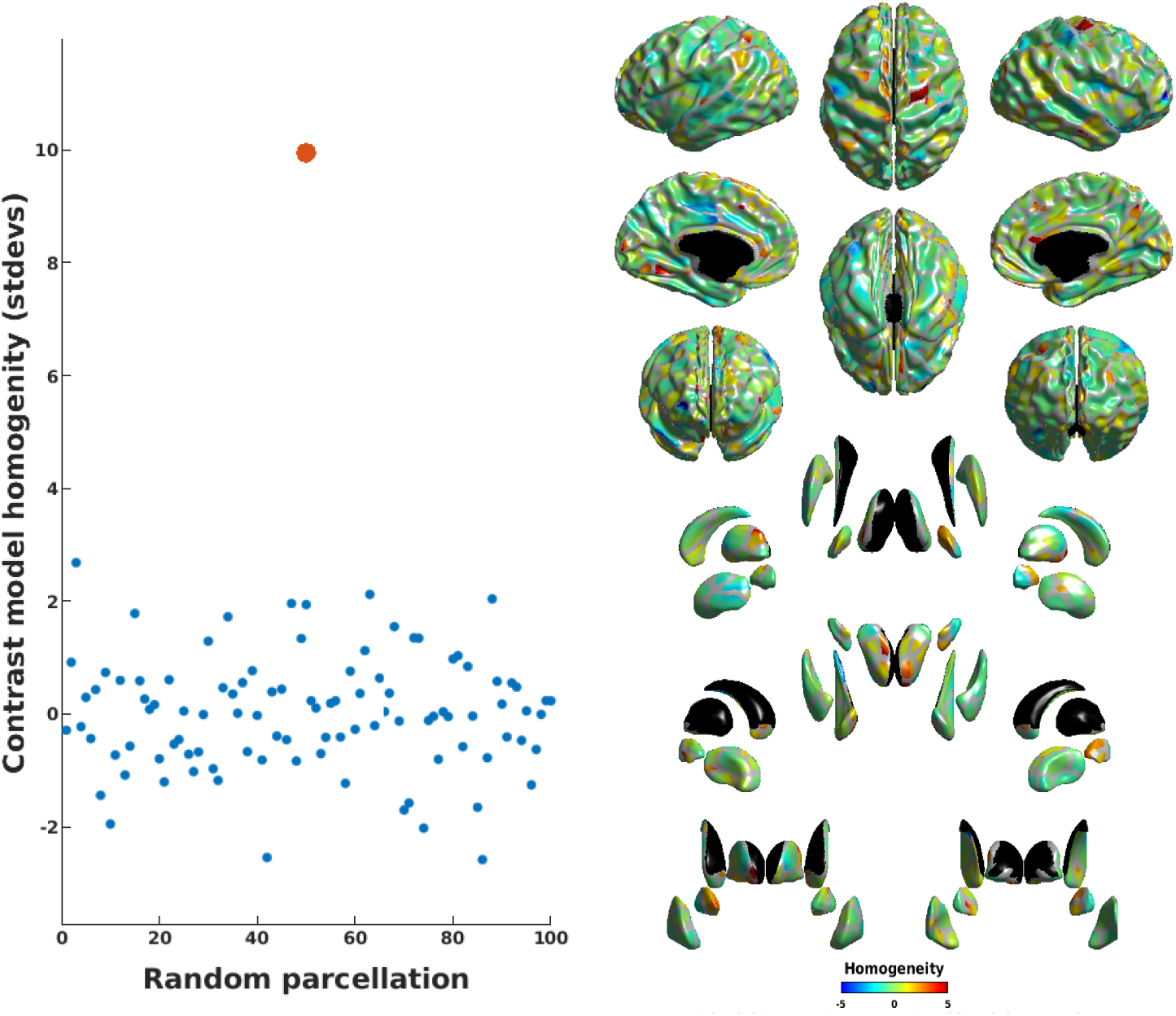
White/gray contrast model complexity homogeneity. The left subfigure shows the comparison of mean within-parcel homogeneity of the white/gray contrast model complexity for the BLeG parcellation (red) vs random parcellations (blue). The BLeG parcellation shows mean within-parcel homogeneity that is almost 10 standard deviations greater than that for the random parcellations. The subfigure on the right shows the pattern of the increased homogeneity in the BLeG parcellation.

## 4. Discussion

Drawing on our recent work using label-based fusion methods to identify the thalamus, caudate, putamen, and globus pallidus, which we then fitted surfaces to, we extended Gordon et al.’s (2014) surface-based rs-fMRI parcellation approach to include also these deep gray-matter structures. We generated a functional parcellation of both the cortical and subcortical surfaces using the life-span data from the Enhanced Nathan Klein Institute - Rockland Sample (Nooner et al., 2012), comprised of data from 590 individuals from 6 to 85 years of age. We call this the Bezgin-Lewis extended Gordon (BLeG) parcellation, and we provide it for the use of the neuroimaging community ^3^.

The cortical portion of our BLeG parcellation, unsurprisingly, is similar to Gordon’s parcellation; the methods are, for the most part, identical, other than the subcortical structures here being represented as surfaces rather than volumes. But there are differences in the cortical parcellations. There are 316 cortical BLeG regions (left 160; right 156). The Gordon parcellation has 356 parcels (178 in each hemisphere). There are a number of possible reasons for our slightly coarser parcellation. First, the age range of the NKI-RS data is 6 to 85 years, whereas the Gordon parcellation was based on data from young adults. Functional connectivity patterns are known to be similar across the life-span, but there are some age-related differences (Han et al., 2018); such differences might essentially blur some of the watershed boundaries. Our sample size was also substantially larger than that used by Gordon et al. (2014). Whereas Gordon et al. used a dataset comprised of data from 120 subjects, and a verification dataset comprised of data from 108 subjects, we used the available NKI-RS data, which constituted 590 subjects. Moreover, the NKI-RS data are from a single run, whereas the Gordon data combines multiple runs. Further, there are substantial differences in the resolution of the data: the Gordon et al. (2014) data having a spatial resolution of either 4×4×4 mm or 3×3×3.5 mm, and a temporal resolution of 2.5 s; the NKI-RS sample was acquired at a spatial resolution of 2×2×2 mm, with a temporal resolution of 1400 ms. It is unclear to what extent those differences should yield a coarser or finer parcellation. But the higher spatial resolution of the NKI-RS data used here compared to the data used by Gordon et al. most likely explains another difference between the BLeG parcellation and the Gordon parcellation. Gordon et al. excluded parcels considered unreliable due to low-SNR (Wig et al., 2014). We saw no evidence to support this type of exclusion for our cortical+subcortical approach with the NKI-RS data; the watershed boundaries in those areas excluded by Gordon et al. appeared no less strong than in regions included by Gordon et al.

The watershed boundaries in the subcortical portion of our BLeG parcellation were also clear. There are 16 thalamic BLeG regions (8 in either hemisphere); 18 putamenal BLeG regions (9 in either hemisphere); 6 pallidal BLeG regions (3 in either hemisphere); and 14 caudatal BLeG regions (left 8; right 6). As per Gordon et al. (2014), but for the subcortical structures, we assessed our BLeG parcellation in comparison to the known anatomy of these structures. As shown by the example of the thalamus in section 3.2, our BLeG subcortical parcels seem to be well-aligned with the consensus anatomy. For the thalamus, there was a direct correspondence between the known anatomy and the BLeG parcels. A more thorough comparison of the BLeG parcellation and cytoarchitectonically defined regions would, of course, be better, but the cytoarchitectonic data are lacking for such a comparison. Nonetheless, there are points of apparent disagreement that are worth consideration. The globus pallidus, for instance, appears to comprise two anatomical structures: the globus pallidus internal and external. The BLeG parcellation instead divides the globus pallidus into three regions. But recent work suggests that the globus pallidus does, in fact, divide into three regions: the external segment, the lateral internal segment, and the medial internal segment (Kita, 2010).

Gordon et al. (2014) also previously showed increased within-parcel homogeneity of functional connectivity patterns for their parcellation compared to spatially randomized variants of it. Our analysis repeats that assessment and extends it to include the subcortical parcels. We verify that there is increased within-parcel homogeneity of functional connectivity patterns for our cortical+subcortical BLeG parcellation, and that the subcortical structures also show this increased within-parcel homogeneity. Further, our analysis adds to that a demonstration that the BLeG functional parcels align better with the patterns of the complexity of the models that best fit the white/gray contrast measures at each vertex than do randomly generated parcellations with the same number of parcels and the same mean parcel size, *i.e.* the BLeG parcellation showed generally greater within-parcel white/gray contrast model complexity homogeneity compared with such randomly generated parcellations. This was also true for both the cortical and subcortical parcels.

This cross-modal demonstration provides a verification of the validity of the BLeG parcellation in that different brain regions mature at different rates, show different patterns of structural and functional connectivity, show sexual dimorphism to differing degrees, etc. (Raz et al., 1997; Goldstein et al., 2001; Allen et al., 2005; Raz et al., 2005; Raz and Rodrigue, 2006; Sowell et al., 2006; Kennedy et al., 2009; Storsve et al., 2014; Østby et al., 2009; Goddings et al., 2014). Thus, we expect that the models that provide the best-fit to the data will be similar within brain regions, and so show high within-parcel model complexity homogeneity. This cross-modal demonstration also provides evidence of the usefulness of the BLeG parcellation for analyses of data derived from other than fMRI.

Additionally, the BLeG parcellation may provide a bridge between studies using samples with different age ranges. As noted by Han et al. (2018), parcellations based on samples from different time periods across the life-span show far more similarities than differences, but the differences preclude direct comparisons. And the changes in morphology over the life-span make a volumetric approach untenable (Anticevic et al., 2008; Klein et al., 2010). Sulci are narrow in children, with blood vessels encased by sulcal walls, whereas in the elderly the sulci are generally more open, with blood vessels adjacent to one or the other sulcal wall. Volumetric blurring in children is thus substantially different than it is in the elderly. Surface-based approaches are less impacted by these differences, as the data are read onto the surface and smoothing is along the 2D manifold. Thus the BLeG parcellation, based on data from 6 to 85 years of life is arguably a good parcellation to use to allow results from different samples to be compared.

But some caveats should be noted. Though there are certainly gains made by using a surface-based approach in terms of registration (Fischl et al., 2007; Lyttelton et al., 2007; Anticevic et al., 2008; Klein et al., 2010), there are also potentially losses. Whereas the cortex is fairly indisputably, at least at the resolution of fMRI, a 2D structure, that is less clear for the deep gray matter. There is a potentially important third dimension to the deep gray-matter structures. The intralaminar nuclei of the thalamus, for instance, are potentially lost in a surface-based analysis. This could, of course, be dealt with by generating surfaces at multiple depths within the deep gray-matter structures; but we have not done this here. There are also deep gray-matter structures that are difficult to fit surfaces to, *e.g.* the hippocampus; and we have not. Likewise, the cerebellum has such fine structure in terms of gray-matter folds that it is exceedingly difficult to perform tissue segmentation on, and to fit surfaces to; and we have not.

In summary, building on Gordon et al.’s (2014) surface-based rs-fMRI parcellation approach, and on Lewis et al.’s (2019) methods for labelling and putting surfaces on the subcortical material, we have extended Gordon et al.’s method to include the deep structures as well as the cortex. We have applied this extended methodology to the life-span data from the Enhanced Nathan Klein Institute - Rockland Sample (Nooner et al., 2012), comprised of data from 590 individuals from 6 to 85 years of age, and generated the Bezgin-Lewis extended Gordon (BLeG) parcellation. We have shown that our BLeG parcellation has much higher within-parcel rs-fMRI connectivity homogeneity than should be expected based on parcel size alone. We have shown that the subcortical parcels of the BLeG parcellation align with the known anatomy of the subcortical structures. And we have shown that within-parcel model complexity homogeneity for gray/white contrast is much higher for the BLeG parcellation than should be expected based on parcel size alone. We provide the BLeG parcellation for the use of the neuroimaging community with the hope that it proves useful.

## 5. Acknowledgments

This research has been supported by grant ANRP-MIRI13-3388 from the Azrieli Neurodevelopmental Research Program in partnership with the Brain Canada Multi-Investigator Research Initiative (to ACE), and by grants from the Canadian Institutes of Health Research (to ACE and to DLC), and grants from the Natural Sciences and Engineering Research Council of Canada (to DLC). It also benefited from computational resources provided by Compute Canada (www.computecanada.ca) and Calcul Quebec (www.calculquebec.ca).

## Footnotes

1 https://github.com/aces/CIVET_Full_Project

2 SurfStat is available at http://www.math.mcgill.ca/keith/surfstat/

3 http://mcin.ca/research/ — *Once the paper is accepted.*

